# Organoid-based single-cell spatiotemporal gene expression landscape of human embryonic development and hematopoiesis

**DOI:** 10.1101/2022.09.02.505700

**Authors:** Yiming Chao, Yang Xiang, Jiashun Xiao, Shihui Zhang, Weizhong Zheng, Xiaomeng Wan, Zhuoxuan Li, Mingze Gao, Gefei Wang, Zhilin Chen, Mo Ebrahimkhani, Can Yang, Angela Ruohao Wu, Pentao Liu, Yuanhua Huang, Ryohichi Sugimura

## Abstract

Single-cell level characterization of embryonic development is a major benchmark of human developmental biology. Spatiotemporal analysis of stem-cell-derived embryos offers conceptual and technical advances in the field. Here, we defined the single-cell spatiotemporal gene expression landscape of human embryonic development with stem-cell-derived organoids. We established the human embryonic organoid (HEMO) from expanded potential stem cells and achieved both embryonic and extraembryonic tissues in the same organoid. Time-series single-cell RNA sequencing paired with single-cell resolution spatial revealed human embryonic development signatures such as extraembryonic placenta, yolk sac hematopoiesis neural crest, blood vessels, and cardiac mesoderm. Hematopoietic tissues eventually predominated HEMO with erythropoiesis, mekagaryopiesis, and myelopoiesis. Cell-cell communication network analysis demonstrated that trophoblast-like tissues supplied WNT signaling in neural crest cells to facilitate maturation and migration. Single-cell resolution spatial transcriptomics defined the yolk sac erythro-megakaryopoietic niche. Vitronectin-integrin signaling, a major contributor to megakaryocyte maturation, was predominant in the yolk sac niche in HEMO and to human fetal samples. Overall, our study advances the spatiotemporal analysis of human embryonic development in stem-cell-derived organoids.

**Highlights:**

- Modeling human embryonic development from stem cells
- Used of both 10X Chromium and 10X Visium to define the gene expression landscape of embryonic development and hematopoiesis
- WNT signaling as a regulator of neural crest maturation and EMT
- VTN-ITGA2B as the main contributor to Mk maturation within the yolk sac erythro-megakaryopoietic niche

**Graphical abstract:** 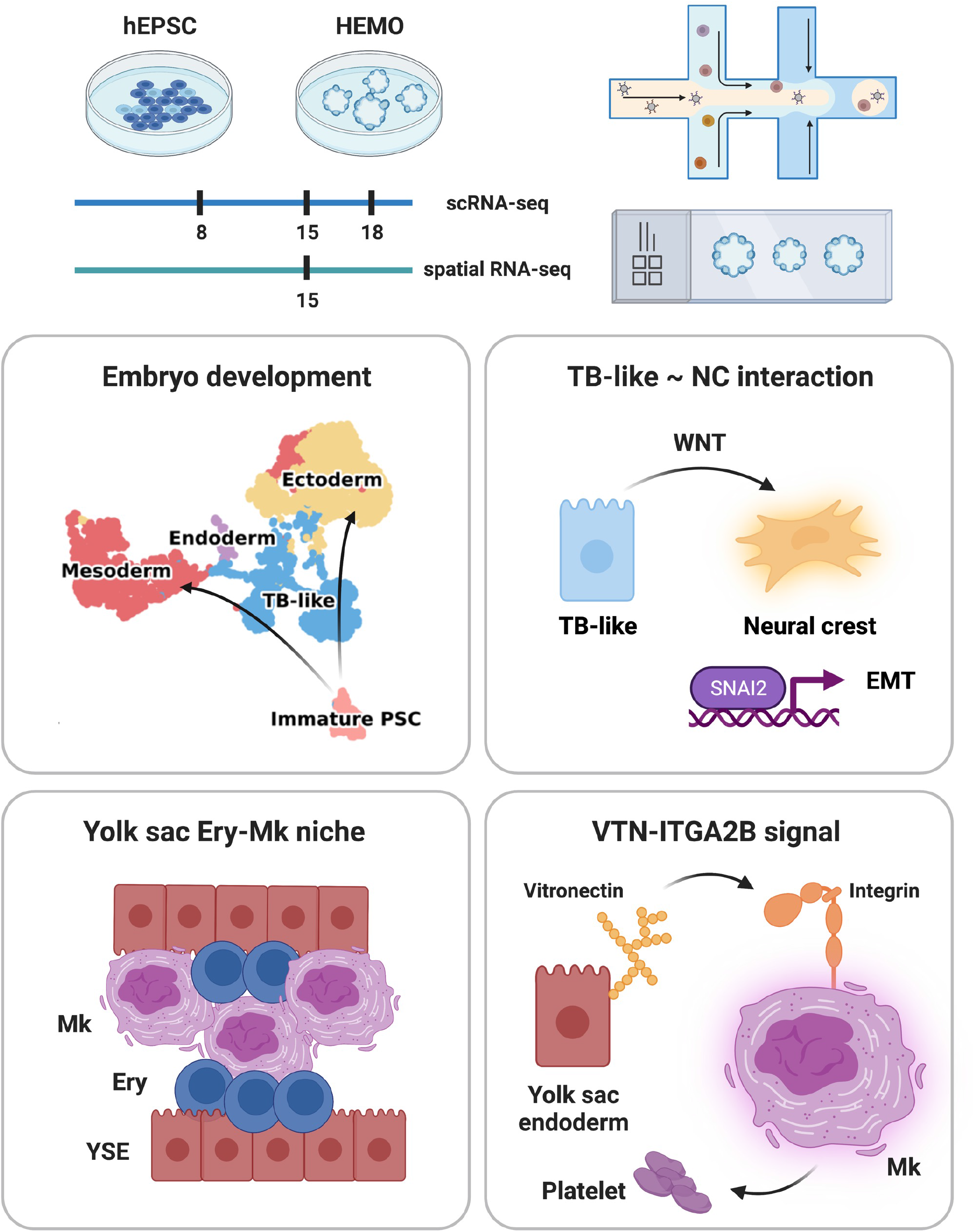

## Introduction

Human embryonic development starts with the partition of both extra- and embryonic tissues. The extraembryonic tissues placenta and yolk sac supply hematopoietic cells and nutrition to support the embryo (Gude et al., 2004; Jollie, 1990). The cell-cell interactions are fundamental mechanisms to differentiate embryonic tissues. It remains unknown if extraembryonic tissues regulate the differentiation of embryonic tissues. The benchmark of differentiation of embryonic tissues is a partition of mesoderm into cardiac mesoderm, vascular endothelium, and hematopoietic cells (Aulehla and Pourquié, 2010). Among these tissues, hematopoietic cells serve as a sight glass of embryonic development with their well-defined lineage trajectory reflecting the stage of embryos (Zeng et al., 2019a). For instance, megakaryopoiesis starts from hematopoietic progenitor cells through megakaryocyte progenitors to megakaryocytes (Noetzli et al., 2019; Tober et al., 2007). Myelopoiesis starts from hematopoietic progenitor cells through common myeloid progenitors to monocytes (Rosenbauer and Tenen, 2007).

Stem-cell-derived embryos reconstitute the developmental process and cell-cell interactions. Previous studies have modeled human and mouse embryonic development from pluripotent stem cells or directly used primary embryos (Aguilera-Castrejon et al., 2021; Alsinet et al., 2022; Anlas and Trivedi, 2021; Atkins et al., 2022; Ávila-González et al., 2022; Decembrini et al., 2020; Diaz-Cuadros et al., 2020; Lancaster et al., 2013; Mackinlay et al., 2021; Matsuda et al., 2020; Minn et al., 2020; Morgani et al., 2018; Saykali et al., 2019; Shahbazi et al., 2016; Simunovic et al., 2022; Tamaoki et al., 2021). Synthetic embryos recapitulated gastrulation and organogenesis by assembling pluripotent stem cells and their extraembryonic derivatives (Amadei et al., 2022; Harrison et al., 2017). The advanced culture method of naïve pluripotent stem cells allows for specification of placenta lineage (Dong et al., 2020; Guo et al., 2021; Io et al., 2021; Liu et al., 2020). Blastoids from naïve pluripotent stem cells model pre-implantation and post-implantation human embryos (Aguilera-Castrejon et al., 2021; Kagawa et al., 2022; Li et al., 2019; Rivron et al., 2018; Tarazi et al., 2022; Vrij et al., 2022; Yu et al., 2021). The recently established expanded potential stem cells (EPSCs) possess the capacity to derive both embryonic and extraembryonic tissues (Gao et al., 2019; Tan et al., 2021; Yang et al., 2017). The use of EPSCs possesses advantages in modeling cell-cell interactions in human embryonic development, especially to examine extraembryonic tissues regulating the differentiation of embryonic tissues.

The advance in single-cell sequencing and increased accessibility of human fetal samples generated various cell atlas of human embryonic development (Bian et al., 2020; Popescu et al., 2019; Rajab et al., 2021; Vento-Tormo et al., 2018; Wang et al., 2021; Zeng et al., 2019a, 2019b). Pairing these datasets with the stem-cell-derived embryos deciphers cell-cell interactions in development. The recent emergence of spatial transcriptomics technology has generated various tissue atlas across various species (Asp et al., 2019; Chen et al., 2021; Fawkner-Corbett et al., 2021; Fink et al., 2022; Mantri et al., 2021; Maynard et al., 2021; Moses and Pachter, 2022; Tower et al., 2022; Zhao et al., 2022). Spatial transcriptomics can faithfully restore the original spatial location, allowing us to unravel the functional niche and explore the cellular signaling interactions. However, one of the technology-related limitations is the limited resolution of cells. Thus, more advanced computational methods are required to achieve single-cell resolution spatial transcriptomics.

We aim to study human embryonic development in stem-cell-derived organoids. Given the technology advance of high throughput sequencing methods, we exploited the time series single-cell (sc)RNA-sequencing and spatial transcriptomics to study cell-cell interactions in development of embryonic tissues.

Here we leveraged an EPSC-derived Human EMbryonic Organoid (HEMO) to model development of embryonic tissues. The time-series single-cell RNA sequencing showed that HEMO follows the development of human embryos with the subsequent emergence of extraembryonic tissues and three germ layers. We highlighted that the WNT signals secreted from trophoblast-like tissue act as a potential mediator for neural crest cell maturation and migration. With 10X Visium spatial transcriptomics and an advanced cell deconvolution method, we defined the yolk sac erythro-megakaryopoietic niche and revealed vitronectin-integrin as the main contributor to megakaryocyte maturation. Collectively, HEMO proposes a model of human embryonic development and hematopoiesis.

## Results

### Time-series scRNA-seq of the human embryonic development and hematopoiesis in stem-cell-derived organoids

To model human embryonic development, we used expanded potential stem cells (EPSCs) which possess both embryonic and extraembryonic differentiation potential (Gao et al., 2019). We established a human embryonic organoid (HEMO) with an adaptation of a Stemdiff hematopoiesis kit in a three-dimensional culture (Fig. 1A). HEMOs followed germ layer patterning and enrichment of hematopoietic lineage with morphogens and cytokines.

**Fig 1.**
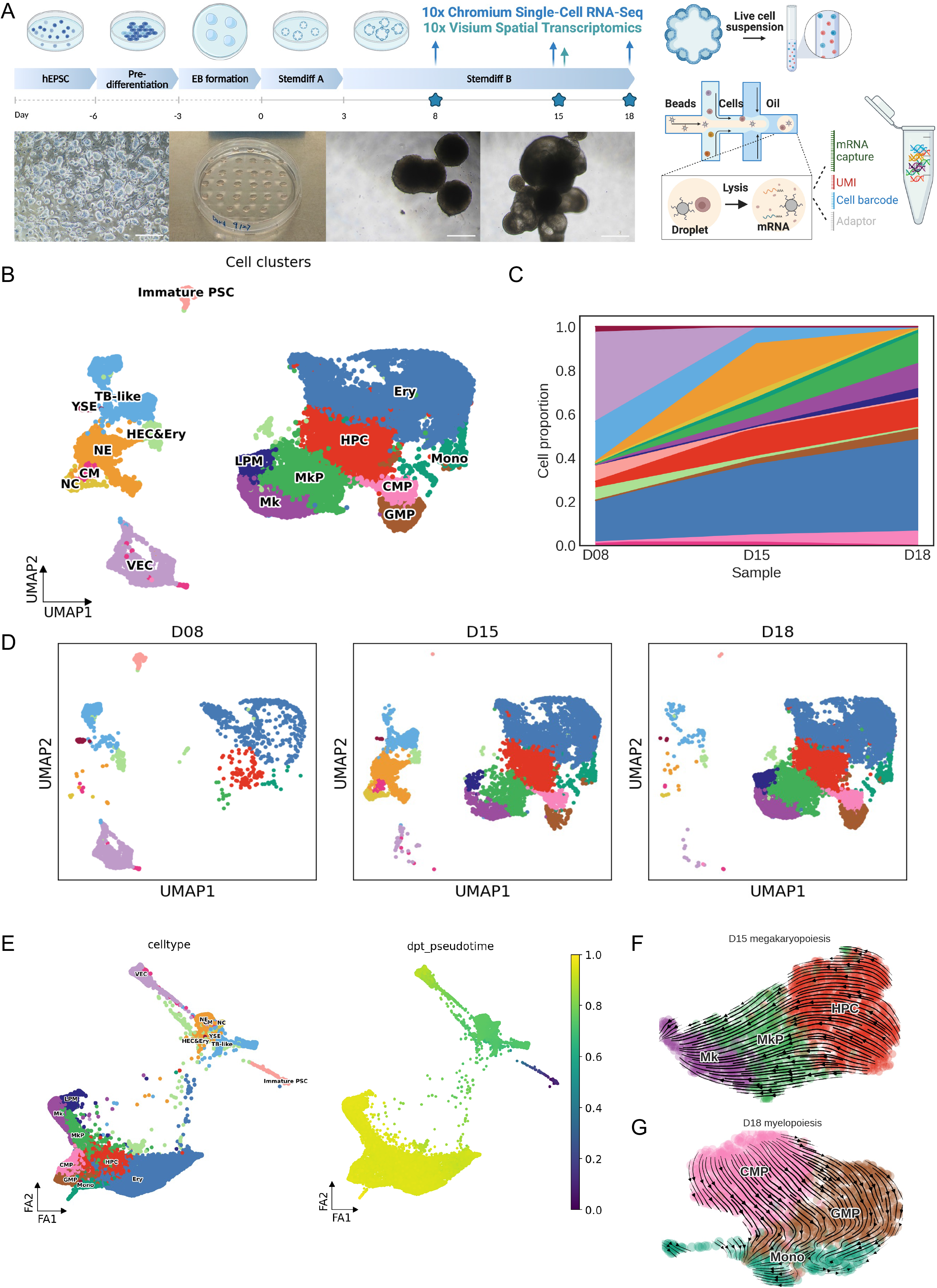
Time-series scRNA-seq of the human embryonic development and hematopoiesis in stem-cell-derived organoids. A) Differentiation protocol of human embryonic organoid (HEMO). Human expanded potential stem cells (EPSCs) were maintained in EPSC medium, followed by pre-differentiation and embryonic body (EB) formation in the hanging drop. Experimental design: HEMOs at D8, D15, and D18 were harvested for 10X Chromium scRNA-seq. D15 HEMOs were also harvested for 10X Visium spatial transcriptomics. The right panel illustrates the sample preparation of 10X Chromium scRNA-seq. The scale bar = 500 um. B) scRNA-seq analysis of time-series HEMO cells (n=22,296) visualized by UMAP. CM: Cardiac mesoderm, CMP: Common myeloid progenitors, Ery: Erythroid cells, GMP: Granulocyte monocyte progenitors, HEC&Ery: Hemogenic endothelial cells and erythroid cells, HPC: Hematopoietic progenitor cells, Immature PSC: Immature pluripotent stem cells, LPM: Lateral plate mesoderm, Mk: Megakaryocytes, MkP: Megakaryocyte progenitors, Mono: Monocytes, NC: Neural crest, NE: Neural ectoderm, TB-like: Trophoblast-like cells, VEC: Vascular endothelial cells, YSE: Yolk sac endoderm. C) Area chart shows cell proportion changes during the differentiation. D) Individual UMAP shows cell populations at D8 (n=3,136), D15 (n=8,364), D18 (n=10,796). E) Diffusion pseudotime analysis of the developmental trajectory of HEMO. Cells are embedded into force-directed graphs. F) RNA velocity analysis of D15 megakaryopoiesis following HPC, MkP, Mk. G) RNA velocity analysis of D18 myelopoiesis following CMP, GMP, and Mono.

To examine if HEMO follows human embryonic development, 10X Chromium single-cell RNA-sequencing (scRNA-seq) picked up three-time points to measure extra-embryonic lineage at D8, three germ layers at D15, and hematopoiesis at D18 (Fig. 1A). We first removed the doublets through DoubletFinder and did batch correction with MNN on three samples (Haghverdi et al., 2018; McGinnis et al., 2019). Among 22,296 cells, we annotated sixteen cell clusters by known marker genes (Fig. 1B; Ext. Fig. 1E), including human development-related lineages like immature pluripotent stem cells (Immature PSC), trophoblast-like cells (TB-like), yolk sac endoderm (YSE), cardiac mesoderm (CM), vascular endothelial cells (VEC), neural ectoderm (NE) and neural crest (NC). We categorized hematopoietic cell clusters into hemogenic endothelial cells and erythroid cells (HEC&Ery), hematopoietic progenitor cells (HPC), erythroid cells (Ery), megakaryocyte progenitors (MkP), megakaryocytes (Mk), common myeloid progenitors (CMP), granulocyte monocyte progenitors (GMP) and monocytes (Mono). We identified the feature of lateral plate mesoderm (LPM), an ancestor of hematopoietic lineage, clustered close to hematopoietic populations.

The extra-embryonic tissues TB-like cells and YSE emerged at D8. The embryonic trunk tissues CM, NE, and NC emerged at D15 (Fig. 1C, D). The hematopoietic population became predominant at D18 (Fig. 1E). To examine the maturation of hematopoietic cells, we adapted RNA velocity (Gao et al., 2022). The megakaryopoiesis started from hematopoietic progenitor cells, through megakaryocyte progenitors, to megakaryocytes, which is consistent with the literature knowledge (Fig. 1F) (Noetzli et al., 2019). The myelopoiesis started from CMP, through GMP, to monocyte, which again emphasized the known process (Rosenbauer and Tenen, 2007) (Fig. 1G). According to the phase portrait showing transcriptional activities, we found that the maturation of ITGA2B mRNAs as an indicator of megakaryopoiesis (Ext. Fig. 1F).

Altogether, HEMO followed a human embryonic development. Time-series scRNA-seq of HEMO revealed the gene expression landscape of human embryonic development and hematopoiesis at the single-cell level.

### HEMO acquires development of endoderm, mesoderm, ectoderm, and trophoblast-like tissues

To examine three germ-layer patterning of HEMO, we further grouped non-hematopoietic clusters into Immature PSC, TB-like, Ectoderm (NE, NC), Mesoderm (VEC, CM) and Endoderm (YSE) (Fig. 2A). These cell populations indicate the patterning of three germ layers and trophoblasts (Fig. 2B). We identified marker genes for Immature PSC (NANOG, SOX17, KLF4, POU5F1), TB-like (CDH1, TFAP2A, EPCAM, KRT7), Ectoderm (SOX3, PAX6, OTX2, NES), Mesoderm (PDGFRA, KDR, BMP4, NCAM1), and Endoderm (FOXA2, AFP, APOA1, GATA4) (Fig. 2C, D). We noted an overall reduction of non-hematopoietic populations and proportion changes when HEMO switched to hematopoietic fate at D18 (Fig. 2E, F). Extraembryonic tissues (TB-like, YSE) emerged first, followed by mesoderm (VEC, CM) then ectoderm (NE, NC). Immature PSCs accounted for a minor population that disappeared by D15 (Ext. Fig. 2A). We then dissected Ectoderm populations which became predominant at D15. We identified embryonic trunk tissues neural crest (NC), neuroectoderm (NE), and cardiac mesoderm (CM) (Fig. 2G). These observations demonstrate that three germ-layer patterning of HEMO followed the development of endoderm, mesoderm, ectoderm and trophoblast-like tissues.

**Fig 2.**
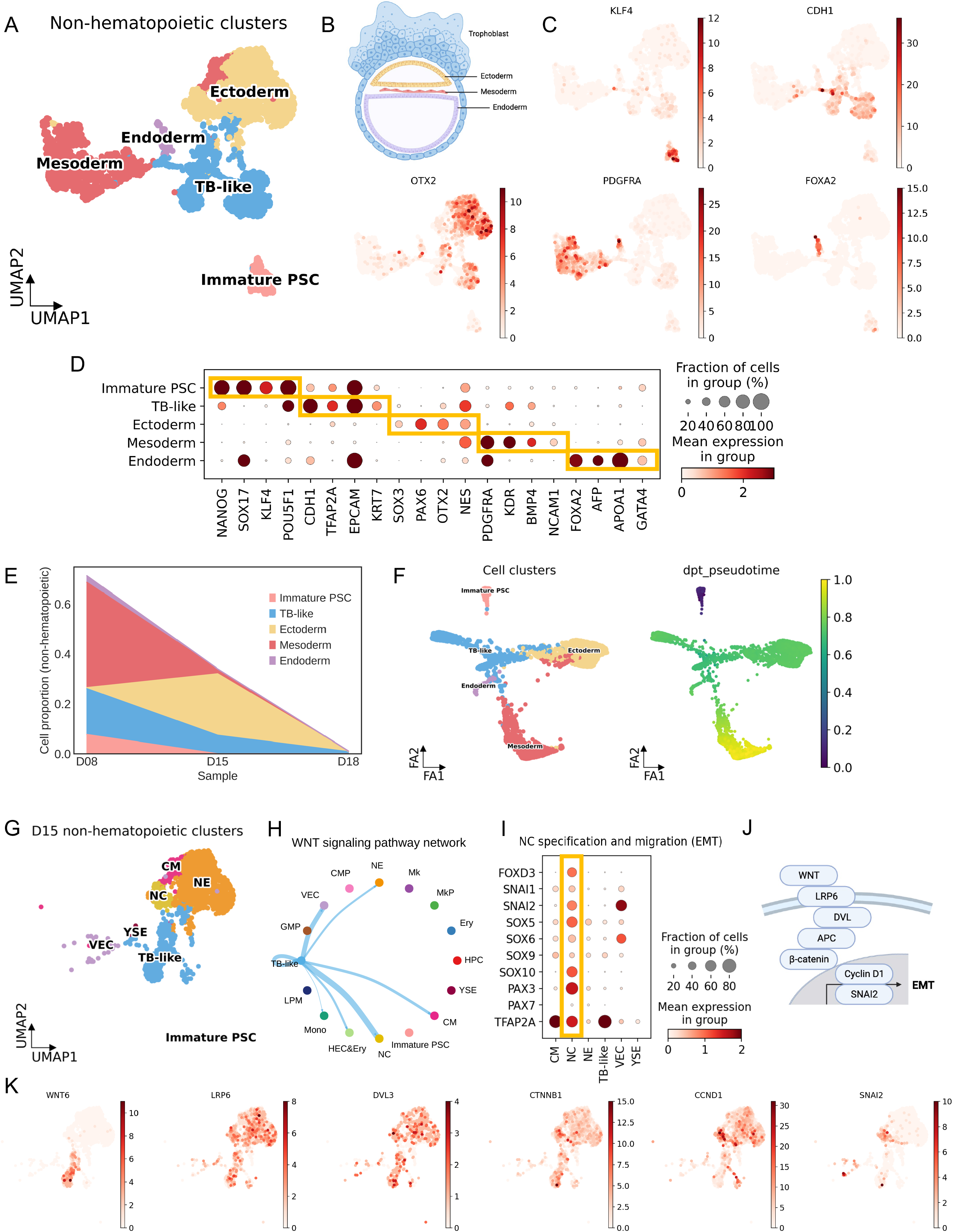
HEMO acquires development of endoderm, mesoderm, ectoderm, and trophoblast-like tissues. A) UMAP analysis of non-hematopoietic cell clusters grouped by Immature PSC, TB-like, Ectoderm (NE, NC), Mesoderm (VEC, CM), Endoderm (YSE) populations (n= 5,203). B) Schematic of early human embryo germ layers, including trophoblast, ectoderm, mesoderm, and endoderm populations. C) Identification of cell populations visualized by UMAP. KLF4, Immature PSC. CDH1, TB-like. OTX2, ectoderm. PDGFRA, mesoderm. FOXA2, endoderm. D) Marker gene expression of each cell cluster. E) Area chart shows cell proportion changes and non-hematopoietic cell number reduction along the differentiation. F) Diffusion pseudotime analysis shows the developmental trajectory of HEMO. Cells are embedded into force-directed graphs. G) UMAP analysis of cell populations in D15 HEMO. H) WNT signaling pathway interaction between D15 HEMO cell populations. Wider lines mean stronger interaction. I) Gene expression pattern of NC specification and migration in D15 non-hematopoietic cells in HEMO. J) Schematic of WNT signaling and involved genes. K) UMAP shows the gene expression of WNT signaling in D15 HEMO.

We then investigated the cell-cell interactions during the development of embryonic trunk tissues by CellChat (Jin et al., 2021). CellChat identified that WNT signaling from TB-like tissues was predominantly connected to neural crest populations (Fig. 2H; Ext. Fig. 2B). The high expression of neural crest specification genes (FOXD3, SOX5, SOX6, SOX9, SOX10, PAX3, PAX7, TFAP2A) and migration genes (SNAI1, SNAI2) (Fig. 2I) suggested the maturation and migration of neural crest cells. We examined if TB-like cells expressed WNT ligands and neural crest expressed mediators of WNT signalings. Indeed, ligands WNT4 and WNT6 were expressed from TB-like cells, whilst receptors LRP6 were expressed in neural crest and neuroectoderm (Fig. 2J, K). WNT mediators DVL, APC, and CTNNB1 lead to the expression of CCND1 and SNAI2, the latter facilitated EMT in neural crest (Fig. 2J, K) (Nusse and Clevers, 2017). These observations suggest a potential function of TB-like tissues in promoting neural crest maturation and migration in the HEMO. We further checked the expression dataset of human fetal spinal cord tissues at Carnegie Stage (CS) 12 and adapted the published annotation (Ext Fig 2C) (Rayon et al., 2021). We found that the neural crest population expressed progenitor markers FOXD3, SOX10 and TFAP2A (Ext Fig 2D). However, they did not express the EMT markers SNAI1 and SNAI2. This suggests that fetal neural crest cells at CS12 are immature, while those of HEMO are at the advanced stage. We note that HEMO is equivalent to CS12 based on the differentiation stage of hematopoietic cells (Ext Fig 2D).

To summarize, HEMO followed a human embryonic development, initiated with the patterning of extraembryonic tissues and neural crest before enrichment of hematopoiesis. TB-like tissue possibly promotes the maturation and migration of neural crest cells via the WNT-driven EMT program.

### Spatial transcriptome reveals cell-cell interactions within the yolk sac hematopoietic niche

To examine the cell-cell interactions in hematopoietic tissues, we conducted spatial transcriptomics on HEMOs at D15 when the diversity of tissue types peaked. To gain single-cell level resolution, we combined 10X Visium with H&E imaging. Capturing nuclei in each Visium spot, we annotated 1-14 cells per spot. The SpatialScope, a framework to quantify the number of a nucleus in each spot, imputed the real cell types based on the paired scRNA-seq dataset (manuscripts in preparation). SpatialScope first did nuclei segmentation to estimate the approximate cell number within each spot. Then, with the paired scRNA-seq data, it assigned the cell type to each nucleus based on a generative model. We achieved approximate single-cell resolution spatial transcriptomics data (Fig. 3B, C). To validate the predicted cell proportion, we applied a standard method RCTD to plot the overall proportion in each spot, which revealed consistency (Ext. Fig. 3A) (Cable et al., 2022).

**Fig 3.**
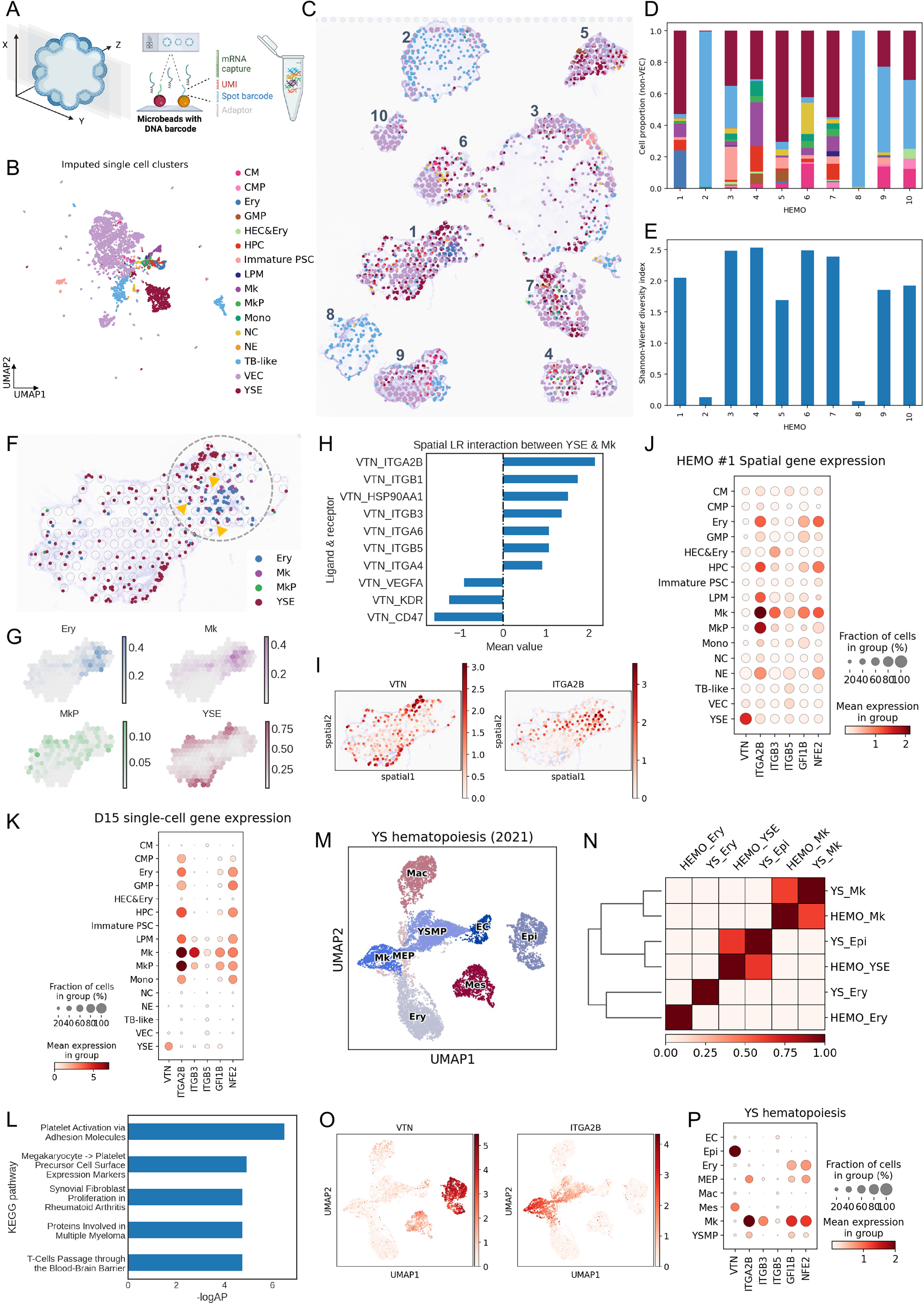
Spatial transcriptome reveals cell-cell interactions of the yolk sac hematopoiesis. A) Illustration of 10X Visium spatial transcriptomics sample preparation on HEMOs. B) UMAP of cell clusters of imputed single-cell level spatial transcriptomic data by SpatialScope. C) Spatial plot shows the HEMOs and single cells imputed by SpatialScope. Spots have been deconvoluted to single-cell level based on nuclei segmentation and transcriptome from scRNA-seq reference. D) Stacked bar plot shows the non-VEC cell composition of each HEMO. E) Shannon diversity index of non-VEC cell types. A larger number indicates more diverse cell types. F) Spatial plot of Ery, Mk, MkP, and YSE population within HEMO #1. The yolk sac erythro-megakaryopoiesis niche is pointed out by a dashed grey circle. Yellow arrows indicate spatial co-localization of YSE, Ery, and Mk in the same spot. G) Cell type deconvolution of HEMO #1 predicted by RCTD. The scale bar indicates the proportion of the specific cell type in each spot. H) Spatial cell-cell interaction analysis conducted by CellPhoneDB. Pairs with a mean value above 0 indicate the activation of the ligand-receptor pairs. Pairs with a mean value below 0 indicate the inhibition of the pairs. Among these, VTN-ITGA2B exhibit the highest enriched score between YSE and Mk. I) Gene expression pattern of VTN and ITGA2B on HEMO #1 in a spatial slice. J) Gene expression of VTN and related downstream targets on imputed spatial data in HEMO #1. K) Gene expression of VTN and related downstream targets on D15 scRNA-seq dataset. L) Bar plot of enriched KEGG pathways based on activated gene targets based on panel I). M) UMAP analysis of cell clusters of in vivo yolk sac hematopoiesis dataset (Wang et al., 2021). Ery, erythrocyte; MK, megakaryocyte; MEP, MK-erythroid progenitor; Mac, macrophage; YSMP, YS-derived myeloid-biased progenitor; EC, endothelial cell; Epi, epithelial cell; Mes, mesenchymal cell. N) Heatmap indicates the correlation between HEMO sample and in vivo yolk sac sample. O) UMAP shows the gene expression of VTN and ITGA2B in the yolk sac dataset. P) Dot plot shows the gene expression of VTN and related downstream targets in the yolk sac dataset.

We then examined the diversity of cell types in each HEMO (Fig. 3C). With the dominant cell type being vascular endothelial cells, we examined the composition of other cell types in each HEMO and calculated the proportion (Fig. 3D). We categorized ten HEMOs based on their cell composition. HEMO #1 contained abundant YSE, erythroid cells, and megakaryocyte lineages, herein called a yolk sac-hematopoietic HEMO (Fig. 3C, D). HEMO #3 predominantly encompassed immature PSCs and TB-like cells, herein called an immature HEMO (Fig. 3C, D). The Shannon diversity index revealed each HEMO-contained diverse cell type except for HEMO #2 and #8, which had predominantly TB-like cells (Fig. 3D, E). These observations indicate that HEMOs develop multiple cell types in 80% of individual organoids.

We further examined cell-cell interactions in the yolk sac-hematopoietic HEMO #1. YSE was mainly located in the margin of HEMO, while erythroid cells and megakaryocyte lineages appeared nearby YSE (Fig. 3F, G). With the single-cell resolution of spatial data by SpatialScope, we detected the co-localization of the YSE, Ery and Mk populations in the same spots, suggesting physical cell-cell interactions. Here, we referred to these regions as the yolk sac erythro-megakaryopoietic niche (Fig. 3 F). We then applied CellPhoneDB to compute the spatial ligand-receptor interaction pairs between four cell types, YSE, Ery, MkP and Mk (Efremova et al., 2020). We observed a strong interactive score between YSE and Mk through vitronectin (VTN) signaling, with VTN-ITGA2B ranked highest (Fig. 3H). The enriched VTN signalings were also supported by another cell type-independent spatial transcriptomics cell interaction tool, SpatialDM (Ext. Fig. 4B) (Li et al., 2022). Among all subtypes, VTN-ITGA2B was the most enriched ligand-receptor pair in HEMO #1. We validated that the VTN and ITGA2B gene expression pattern overlapped with the YSE and Mk areas with RCTD (Fig. 3I, Ext. Fig. 4A). With the single-cell resolution spatial data, we found that VTN was exclusively expressed in YSE, while ITGA2B was expressed in the whole erythro-megakaryopoietic populations (HPC, Ery, MkP, Mk), with the highest expression level in Mk (Fig. 3J). This was also consistent with our RNA velocity analysis that ITGA2B mRNA maturation followed megakaryopoiesis from HPC, MkP to Mk (Ext. Fig. 1F). We also observed high expression of other integrin subtypes ITGB3 and ITGB5 as other receptors of vitronectin known to facilitate hematopoietic development from human pluripotent stem cells (Fig. 3J) (Shen et al., 2021). Notably, GFI1B and NFE2, involved in ITGB3 signaling, were also expressed in Mk, supporting previous studies (Beauchemin et al., 2017; Shiraga et al., 1999). Consistently, we detected expression profiles of vitronectin-integrin genes both in our paired D15 scRNA-seq dataset and spatial transcriptomics (Fig. 3K). The KEGG pathway enrichment analysis revealed platelet activation and megakaryocyte to platelet formation ranked highest, suggesting the role of vitronectin-integrin genes in megakaryocyte maturation and platelet functions (Fig. 3L). These are consistent with the previous literature that VTN-ITGA2B is responsible for platelet aggregation (Asch and Podack, 1990). Hence, we suggest that vitronectin is predominantly expressed by yolk sac endodermal cells, and the integrin pathway promotes megakaryopoiesis in HEMOs.

Finally, we compared HEMOs with the scRNA-seq dataset of human fetal tissue to examine the physiological relevance (Wang et al., 2021). We re-analyzed the dataset of the human fetal yolk sac, herein called ‘YS hematopoiesis’, based on the provided marker list from the reference (Fig. 3M). According to the correlation test, the gene expression pattern of yolk sac erythro-megakaryopoiesis within HEMO resembled the one in the YS hematopoiesis (Fig. 3N). In the YS hematopoiesis dataset, VTN was mainly expressed in yolk sac epithelial cells, and ITGA2B was mainly expressed in megakaryocytes (Fig. 3O). Both YS hematopoiesis and HEMO achieved consistently similar expression patterns of vitronectin, integrins, GFI1B, and NFE2 (Fig. 3P YS hematopoiesis vs Fig. 3K HEMO). Thus, we conclude that HEMOs could mimic human embryonic hematopoiesis in the yolk sac, suggesting vitronectin-integrin signaling as a major contributor to megakaryopoiesis.

## Discussion

The stem-cell-derived embryos have advanced understanding of human embryonic development. This offers a platform to perturb and engineer genes important for human development as well as disease modeling. We leverage the expanded potential stem cells (EPSCs) to generate both extra- and embryonic tissues in the same organoids without the need for separate specification of extraembryonic tissues and assembly. This allows us to explore human embryonic development in an easy operative system. By adopting the defined medium in the commercial kits, we enhanced the reproducibility of the techniques between different labs.

The generation of multiple tissues in each organoid is a hallmark of stem-cell-derived embryos. The current caveat of scRNA-seq is that they cannot distinguish whether each organoid is predominated by a single cell type and a mixture of different organoids, or whether each organoid is genuinely multilineage. Here we leverage spatial transcriptomics and revealed the latter to be the case that HEMO generated multilineage in each organoid, supporting the stem-cell-derived embryos. Combined with time-series scRNA-seq and spatial transcriptomics, we defined the gene expression landscape of human embryonic development and hematopoiesis. HEMO follows a human embryonic development with the acquisition of endoderm, mesoderm, ectoderm and trophoblast-like tissues. We reconstructed cell-cell interactions with ligand-receptor pairs orthogonally with both scRNA-seq and spatial transcriptomics. Our observation that WNT signals from trophoblasts support the maturation of neural crest cells highlights the future of synthetic embryo biology. Although its physiological relevance in real embryos remains to be discovered in future work, the possibility that we could enhance the maturation of certain tissues by perturbation is intriguing. Indeed, compare to the human CS12 in vivo spinal cord dataset, neural crest cells in HEMO were even more mature and functional, as indicated by maturation markers and the EMT program.

The possible application of HEMO is to engineer hematopoietic and immune cells. The contribution of extraembryonic tissue yolk sac in megakaryopoiesis through vitronectin-integrin axis holds a promise to generate platelets from stem cells for cell therapies. The generation of monocytes and macrophages could exploit further engineering to target cancers and infectious diseases. The spatial understanding of the yolk sac hematopoietic niche and leveraging their factors, promotes the generation of immune cells from stem cells.

In sum, HEMO has illustrated the potential to explore functional regulation during human embryonic development and hematopoiesis. Genetically perturbing the process of development in human stem-cell-derived organoids is one of the major applications of HEMO in the research of developmental biology, hematology and immunology. (Sun et al., 2021). HEMO also serves as a source of immune cell engineering for cell therapies.

## Method

### HEMO differentiation

#### hEPSCs maintenance and pre-differentiation

hEPSCs were maintained and cultured in EPSC medium with medium change every other day. Before EB formation, cells were first pre-differentiated in KSR medium (DMEM/F12+10% KSR) for 3 days.

#### EB formation in hanging drop

Remove KSR medium and wash cells with PBS. Digest cells with 500mL 0.05% Trypsin at 37 °C for 3 minutes but no more than 7 minutes. Remove the Trypsin and add 2mL KSR medium and harvest cells. Handle cells gently and avoid harsh pipetting. Centrifuge the cells under 300g for 3 minutes. Remove the supernatant carefully. Add 1mL KSR medium and 1uL Y27632 to resuspend the cells. Add 4k cells for each 25uL hanging drop on the cap of 10 cm petri dish. Make 30-40 drops for each cap and fill the dish with PBS to keep moist. Invert the cap gently and slightly to cover the dish. Label and keep the dishes at 37 °C for 3 days.

#### EB collection and differentiation

Collect all EB with PBS washing. Centrifuge the EB under 100g for 1 min. Remove the supernatant carefully. Add 1mL medium A (STEMdiff Hematopoietic Kit, catalog no. 05310) and transfer to non-adherent 24-well plate. Label the date as Day 0. Culture with 1mL STEMdiff medium A for 3 days. Half-change the medium on Day 2. Culture with 1mL STEMdiff medium B for the following days.

### 10x Chromium scRNA-seq sample preparation and sequencing

#### Preparation of single-cell suspensions

Organoids were harvested at D8, D15, and D18 since EB formation in hanging drop. Organoids were washed with PBS, followed by mechanical chopping with scissors 20-30 times. The tissues were digested in 500 μL Accumax at 37 °C for 10-15 minutes and terminated with 500 μL PBS with 2% FBS. Cells were collected through the 40 μm cell filter. The cell suspension was centrifuged at 500 × g for 5 min and resuspended in FACS sorting buffer (1 × PBS with 2% FBS) for subsequent staining. Cell concentration was adjusted to around 3k cells per μL by counting with a hemocytometer.

#### Flow cytometry for scRNA-seq

Cells were stained in FACS sorting buffer with DAPI (BD Biosciences, catalog no. 564907) in 1:100 for 5 min at 4 °C. Cells were sorted by BD Influx flow cytometry equipment in CPOS at HKUMed. Cells were gated to exclude dead cells and doublets and collected in a chilled single-cell suspension medium (1x PBS with 0.04% BSA) for scRNA-seq library construction. Cell concentration was adjusted to around 500-1000 cells per μL counted with a hemocytometer.

#### scRNA-seq library preparation and sequencing

Single-cell encapsulation, library preparation, and sequencing were done at The University of Hong Kong, LKS Faculty of Medicine, Centre for PanorOmic Sciences (CPOS), Genomics Core. Single-cell encapsulation and cDNA libraries were prepared by Chromium Next GEM Single Cell 3’ Reagent Kit v3.1 and Chromium Next GEM Chip G Single Cell Kit. Around 8-16k live single cells of size 30 μm or smaller and of good viability were encapsulated, followed by reverse transcription and library preparation to harvest a pool of cDNA libraries. Libraries were sequenced using Illumina Novaseq 6000 for Pair-End 151bp sequencing. Individual samples had an average throughput of 165.5 Gb.

### 10x Visium sample preparation and sequencing

#### Frozen sample preparation

Fresh HEMOs were collected on day 15 of differentiation. Washed the HEMOs twice with PBS and transferred the HEMOs into a disposable plastic cryo-mold (Sakura, catalog no. 25608-922), located in the center. Added O.C.T. (Sakura, catalog no. 4583) to immerse the HEMOs fully. Placed the cryo-mold onto the dry ice box until O.C.T. froze (at least 10 min), then stored the cryo-mold immediately at -80C.

#### Reagents and slides preparation

We used the Visium Spatial Gene Expression Starter Kit (10 x Genomics, catalog no. PN-1000200). The composition is the Visium Spatial Tissue Optimization Slide & Reagent Kits (10 x Genomics, catalog no. PN-1000193), Visium Spatial Gene Expression Slide & Reagent Kits (10 x Genomics, catalog no. PN-1000184), and Visium Accessory Kit (10 x Genomics, catalog no. PN-100194).

#### RIN (RNA integrity) detection

Cryosections were cut at 10-μm thickness. Cryosections were flattened out by gently touching the surrounding O.C.T. with cryostat brushes diagonally. We collected at least 25 slices of 10-mm-thick cryosections. RNA was extracted and then analyzed by NanoDrop2000 (Thermo Scientific, catalog no. ND2000CLAPTOP). RIN values were acquired from RNA 6000 Nano). The qualified RIN value should be over 7.0.

#### Tissue Optimization

Tissue Optimization kits were used to optimize the tissue permeabilization time. Tissue cryosections were attached, fixed, stained, and permeabilized for different lengths of time. Fluorescent nucleotide was used as an indicator of permeabilization. The permeabilization time was determined by the highest brightness and lowest diffusion of permeabilized samples under the fluorescent microscope. The optimal permeabilization ensured the release of mRNA and minimized the diffusion of mRNA. In this study, tissue permeabilization time ranged from 6 to 12 minutes.

#### Staining and Tissue imaging

HEMO samples on day 15 of differentiation were embedded in O.C.T. and snap-frozen by a dry ice box. Cryosections were cut at the 10-μm thickness and attached to the GEX slides. The slides were placed on the Veriti™ 96-Well Fast Thermal Cycler (Applied Biosystems, catalog no. 4375305) and incubated for 1 minute at 37C. After incubation, slides were transferred into the pre-chilled methanol at -20C for the fixation, then proceeded to H&E staining. Samples were incubated in isopropanol (catalog no. I9516-25ML) for 1 minute, in Hematoxylin (catalog no. MHS16-500ML) for 7 minutes, after 15 times of quick washing, bluing buffer (catalog no. CS70230-2) for 2 minutes, and Eosin Mix (catalog no. HT110216-500ML) for 1 minute. Slides were incubated onto the thermocycler for 5 minutes at 37C. The brightfield image was obtained by NanoZoomer (Hamamatsu, catalog no. C13239-01) at 20 x resolution.

#### cDNA synthesis and Second strand synthesis

For cDNA synthesis, the stained slides were placed in the Visium Slide Cassette (provided in the Visium kit). The Permeabilization Enzyme (provided in the Visium kit) was added to the wells from the slide cassette. The slide cassette was incubated for 6 minutes at 37C. After washing the wells with 0.1 x SSC (Millipore Sigma, catalog no. S66391L) buffer, we added RT Master Mix (provided in the Visium kit) and initiated reverse transcription for 45 minutes at 53C. After the removal of RT Master Mix, 0.08M KOH (Millipore Sigma, catalog no. P4494-50ML) solutions were added to the well and the slides were incubated for 5 minutes at room temperature. After washing by Buffer EB (QIAGEN, catalog no. 19086), Second Strand Mix (provided in the Visium kit) was added to each well and the slides were incubated for 15 minutes at 65C. The slides were then washed by Buffer EB, and 0.08M KOH was added to the well for 10 minutes at room temperature. Samples from each well were collected and transferred to an 8-tube strip containing Tris solution (Thermo Fisher Scientific, catalog no. AM9850G).

#### cDNA amplification

cDNA cycler number was determined by qPCR (Takara Bio, catalog no. RR820A). The Cq Value was determined to be 25% of the peak fluorescence value. With Amp Mix (provided in the Visium kit) and cDNA Primers (provided in the Visium kit), cDNA was amplified with the determined cycle according to the general PCR protocol.

#### Spatial Gene Expression Library Construction

Library construction was carried out with a library construction kit (10 x Genomics, catalog no. PN-1000190) according to the manufacturer’s protocol. cDNA samples were processed by fragmentation, end-repair & A-tailing, SPRI (Solid Phase Reversible Immobilization) selection, adaptor ligation, and index PCR. The libraries were flanked with P5 and P7 sequences.

#### Sequencing

The Visium libraries consisted of standard Illumina constructs flanked with P5/P7. TruSeq Read 1 was used to sequence the spatial barcode and UMI. TruSeq Read 2 was used to sequence the cDNA insert. Sequence depth ranged from 28.4 Gb to 36.3 Gb.

### Data analysis and statistics

#### scRNA-seq data processing and cell filtering

Sequencing data from 10X Genomics was processed with CellRanger software (version 6.0.0) with GRCh38 human reference genome. Cell by gene count matrixes was loaded to the Seurat package (version 4.1.0) in R (Hao et al., 2021). Cells with fewer than 1,000 detected genes and with more than 15% mitochondrial gene expression were removed. Genes that were expressed in fewer than 3 cells were also removed. Doublets were detected with the package called DoubletFinder (version 2.0.3) for an individual sample with fine-tuned parameters (McGinnis et al., 2019).

Data integration, clustering, and annotation of scRNA-seq data. After all quality control and doublets removal, the scRNA-seq datasets were integrated with MNN (Haghverdi et al., 2018). High variable genes (HVGs) were identified and used for principal component analysis (PCA) for dimension reduction and clustering. Markers for each cluster were identified with the FindAllMarkers function and used for cluster annotation. For non-hematopoietic cell clusters, Immature PSC, TB-like, NE, NC, VEC, CM and YSE were re-clustered into different germ-layer populations.

#### Cell interaction within scRNA-seq data

Cell interaction was analyzed by CellChat (version 1.4.0) (Jin et al., 2021). We followed the standard workflow, loaded the gene expression matrix of the individual sample to CellChat package and calculated the ligand-receptor pair probability. Core functions including computeCommunProb, computeCommunProbPathway and aggregateNet were run following the pipeline. The results were shown in circle graphs.

#### Diffusion pseudotime analysis of scRNA-seq data

The connectivity of the single-cell clusters was first quantified and generated as a partition-based graph abstraction graph (PAGA graph) in Scanpy (version 1.9.1) (Wolf et al., 2019, 2018). For estimating pseudotime, an extended version of diffusion pseudotime (DPT) was applied (Haghverdi et al., 2016). Cells are embedded into force-directed graphs.

#### RNA velocity of scRNA-seq data

Spliced and unspliced RNA were counted by Velocyto (version 0.17.17) and merged with Scanpy adata using loompy function (La Manno et al., 2018). RNA velocity was analyzed by UniTVelo (version 0.1.6) (Gao et al., 2022). UniTVelo proposed a unified latent time across the transcriptome, allowing the incorporation of dynamic genes with weak kinetic information. After subsetting D15 ‘HPC’, ‘Mk’, ‘MkP’ cell clusters for megakaryopoiesis and D18 ‘CMP’, ‘GMP’, ‘Mono’ cell clusters for myelopoiesis, spliced/unspliced information was loaded to UniTVelo for RNA velocity analysis in unified-time mode. The phase portraits and transcriptional activities of interested genes were visualized in scatter plot.

#### 10x Visium spatial transcriptomics processing

Sequencing data and imaging tiff files of individual 10x Visium slides were processed with SpaceRanger software (version 1.3.1) with GRCh38 human reference genome. Count matrixes were loaded to Scanpy package (version 4.1.0) in Python. Spots were selected based on RNA counts and mitochondrial gene percentage. After spot selection, HVGs were identified and used for principal component analysis PCA for dimension reduction and clustering. The Scanpy adata objects were used for downstream analysis individually.

#### Spatial transcriptomics spot deconvolution

Two packages were used for spatial transcriptomics spot deconvolution. Spatialscope method was first implemented (manuscripts in preparation). It is a framework to quantify a number of a nucleus in each spot, and then impute the real cell types based on the paired scRNA-seq dataset. SpatialScope first did nuclei segmentation to estimate the approximate cell number within each spot. Then, with the paired scRNA-seq data, it assigned the cell type to each nuclei-based cell based on a generative model. RCTD package (version 2.0.0) was also used to validate the cell proportion (Cable et al., 2022). It first takes the single-cell reference and predicts the proportion of each cell type in an individual 10x Visium spot. The predicted weighted matrix was used for visualization in the scatter plot.

#### Cell interaction within spatial transcriptomics data

Two different packages were used for spatial cell-cell interaction analysis. After single-cell resolution spatial data, CellPhoneDB within Squidpy toolbox was first used to predict cellular ligand-receptor interactions (Efremova et al., 2020; Palla et al., 2022). The resulting enriched or depleted ligand-receptor pairs were shown in the bar plot. Another tool SpatialDM was also used for predicting local signal interactions at a spatial level (Li et al., 2022). Spot gene count matrix, predicted cell proportion, and spatial coordinate of each fragment were loaded to SpatialDM. It first identified the significant interacting ligand-receptor pairs. Then it identified local spots for each interaction. After pair selection and spot identification, the interactions were visualized in the scatter plots.

#### KEGG pathway enrichment analysis

KEGG pathway analysis was conducted with GSEApy package (https://github.com/zqfang/GSEApy). The gene list of activated targets of VTN was loaded into the package and enriched to the pathway items. The organism was selected as ‘human’. The gene sets were chosen as ‘Elsevier_Pathway_Collection’. The cut-off value was set at 0.5. The enriched items were shown in the bar plot.

## Data availability

The scRNA-seq and spatial transcriptomics data reported in this study have been deposited in NCBI with the accession number PRJNA855311.

## Code availability

Scripts for analyzing scRNA-seq and spatial transcriptomics have been deposited on GitHub: https://github.com/CHAOYiming/HEMO_analysis.

## Acknowledgment

We are thankful to CPOS at HKUMed for technical assistance in scRNAseq, and FACS. We are thankful to Cheryl Tam for operating the cell sorting and Helen Lockey for English editing. We appreciate Danny Chan, Martin Cheung, Lequan Yu, Joshua Ho, Yuanhua Huang’s lab members for discussion and members of the Centre for Translational Stem Cell Biology for discussion and administrative support. The study is supported by Platform for Technology Fund, RGC ECS 27109921, Seed Fund from the School of Biomedical Sciences, and InnoHK Centre for Translational Stem Cell Biology.

## Author contributions

Y.C., Y.X., and R.S. conceived the study. Y.C. conducted informatics analysis with the guidance of Y.H. and R.S. Y.X. generated organoids with the guidance of P.T. and R.S. J.X., S.Z., W.Z., X.W., Z.L., M.G., G.W., and Z.C. contributed to the informatics analysis. C.Y. and A.R.W. supervised the spatial transcriptomics analysis. M.E. made a critical reading of the study. Y.C., Y.X., and R.S. wrote the manuscript.

## Extended figures

**Ext Fig 1.**
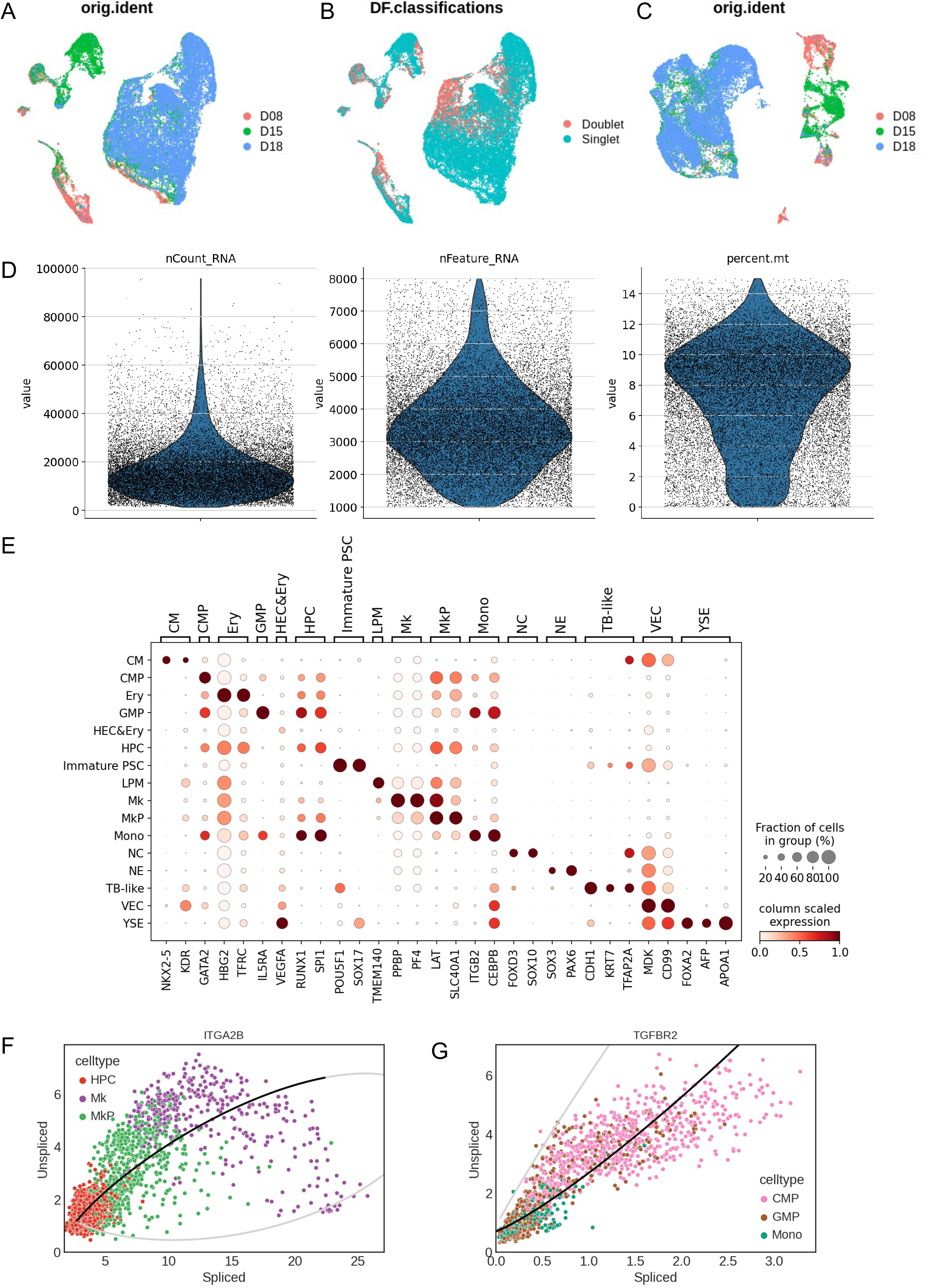
Time-series scRNA-seq of the human embryonic development and hematopoiesis in stem-cell-derived organoids. A) UMAP shows the cell number of three samples. B) Detected doublets and singlets in all samples based on DoubletFinder package. C)Samples after batch correction with MNN package, which are the basis of all downstream analysis. D) Violin plots show the total number of RNA counts, total gene counts, and mitochondrial gene expression percentage of each cell within three samples. E) Marker gene expression of each cell population. F) Scatter plots show expression and splicing/unsplicing level of top gene ITGA2B at D15 megakaryopoiesis. G) Scatter plots show expression and splicing/unsplicing level of top gene TGFBR2 at D18 myelopoiesis.

**Ext Fig 2.**
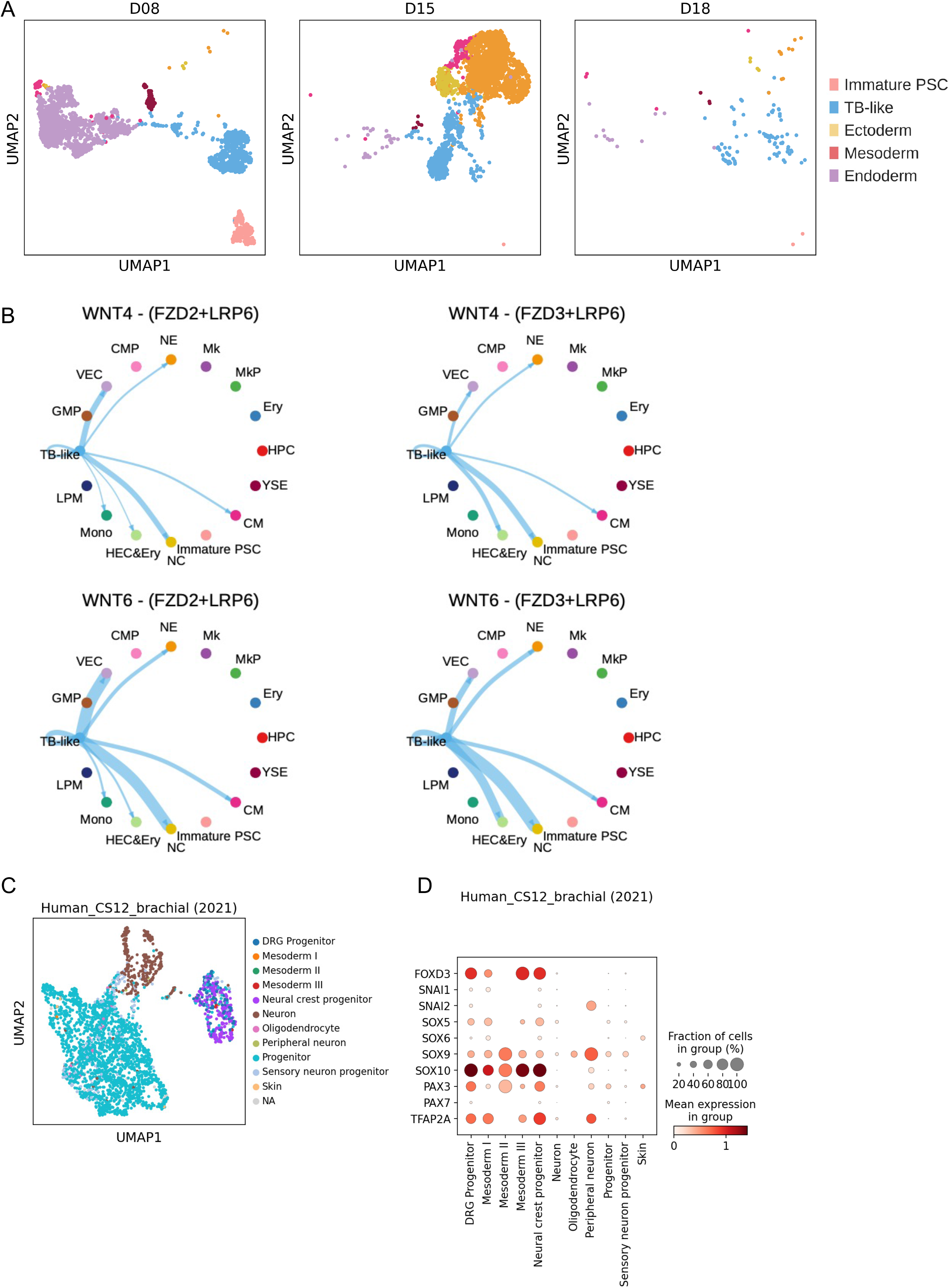
HEMO acquires development of endoderm, mesoderm, ectoderm, and trophoblast-like tissues. A) Individual UMAP of non-hematopoietic populations at D8 (n= 2,236), D15 (n=2,845), D18 (n=122). B) Circle plots show the signaling pathway network of WNT4 and WNT6 subtypes.

**Ext Fig 3.**
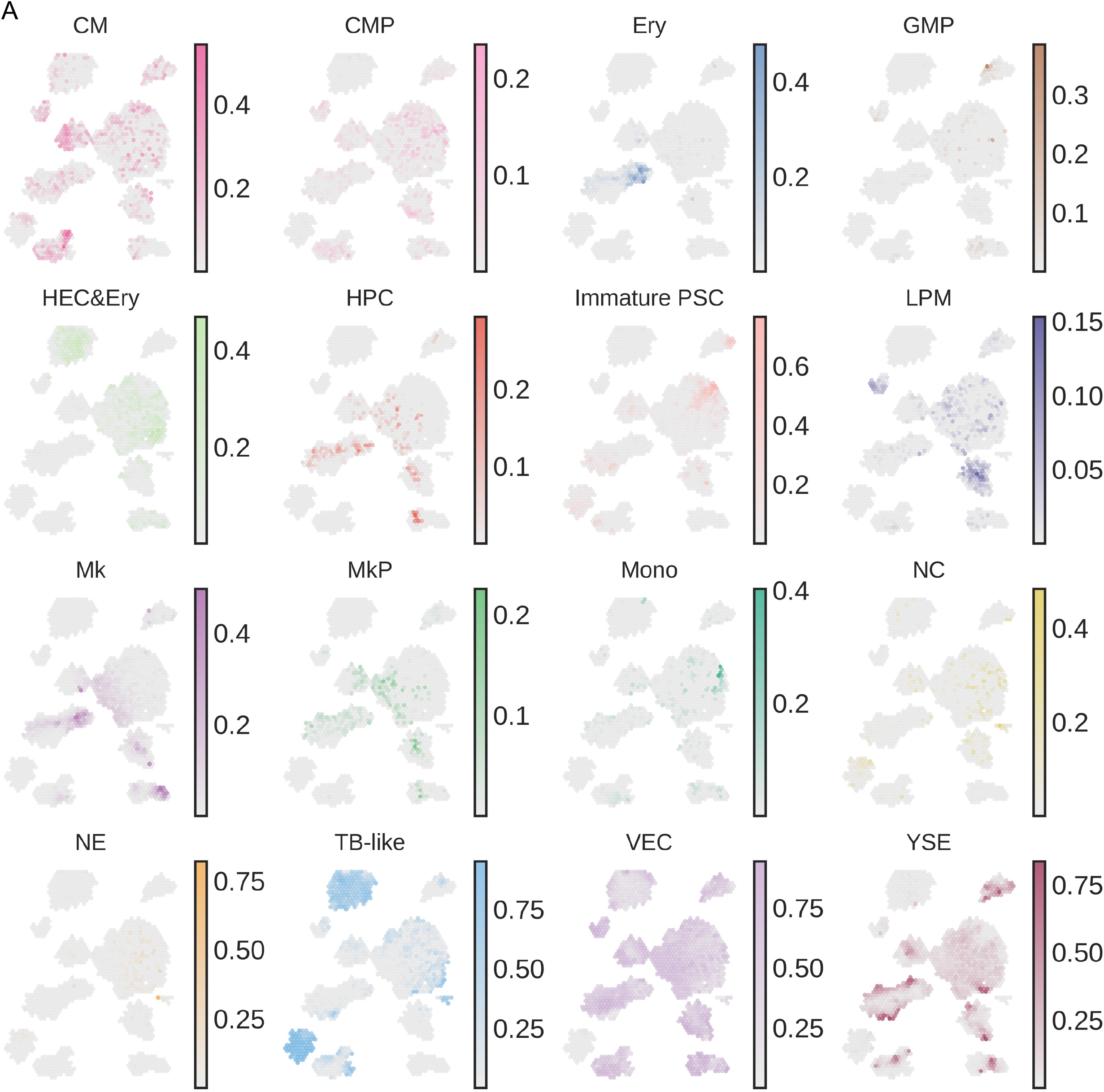
Spatial transcriptome reveals cell-cell interactions of the yolk sac hematopoiesis. A) Scatter plot visualizes cell type deconvolution on HEMO tissues predicted by RCTD. The scale bar indicates the proportion of the specific cell type in each spot.

**Ext Fig 4.**
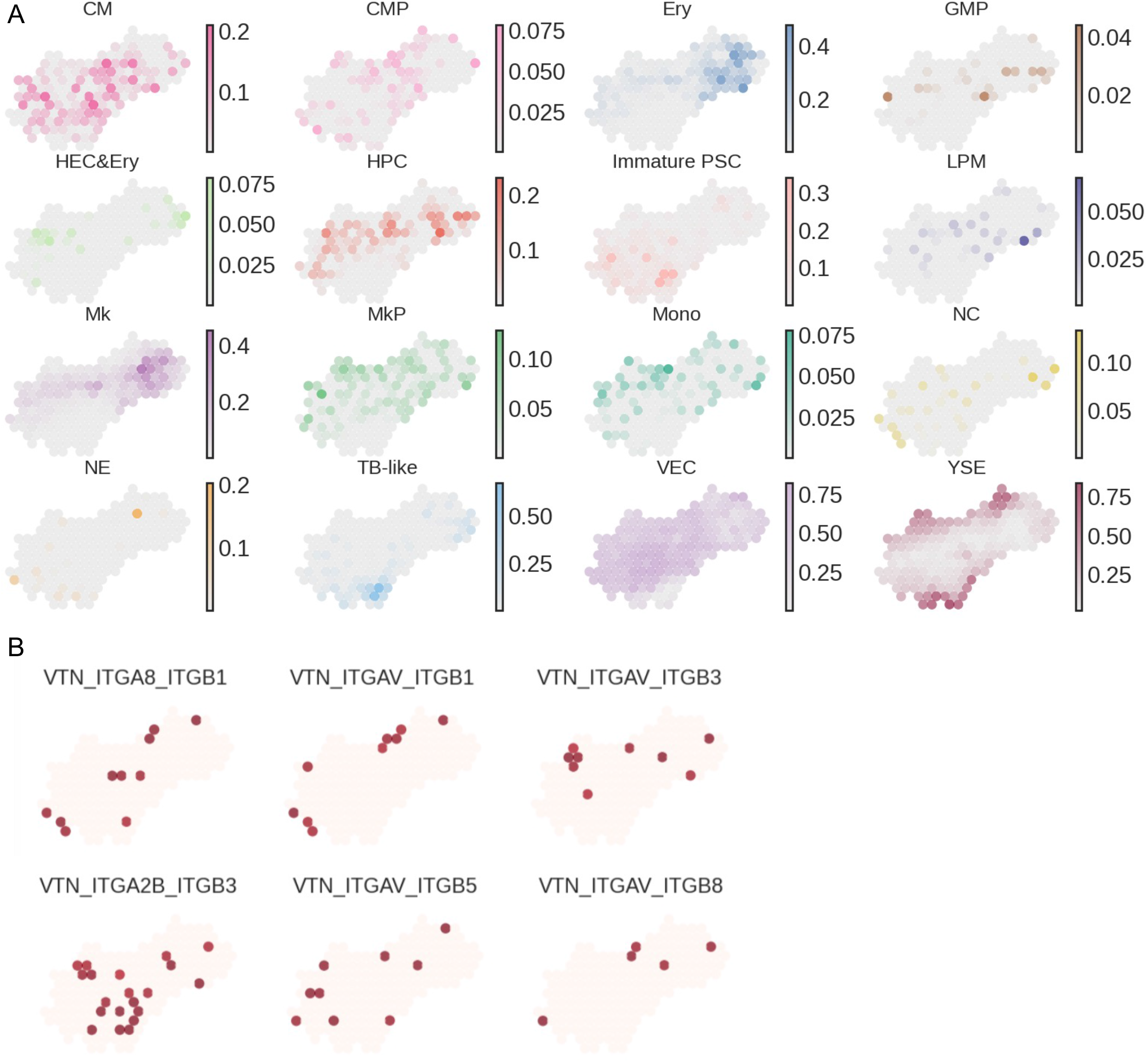
Spatial transcriptome reveals cell-cell interactions of the yolk sac hematopoiesis. A) Scatter plot visualizes cell type deconvolution on HEMO #1 predicted by RCTD. The scale bar indicates the proportion of the specific cell type in each spot. B) Scatter plots of spatial ligand-receptor subtypes of VTN signalings selected by SpatialDM.

**Table 1.**
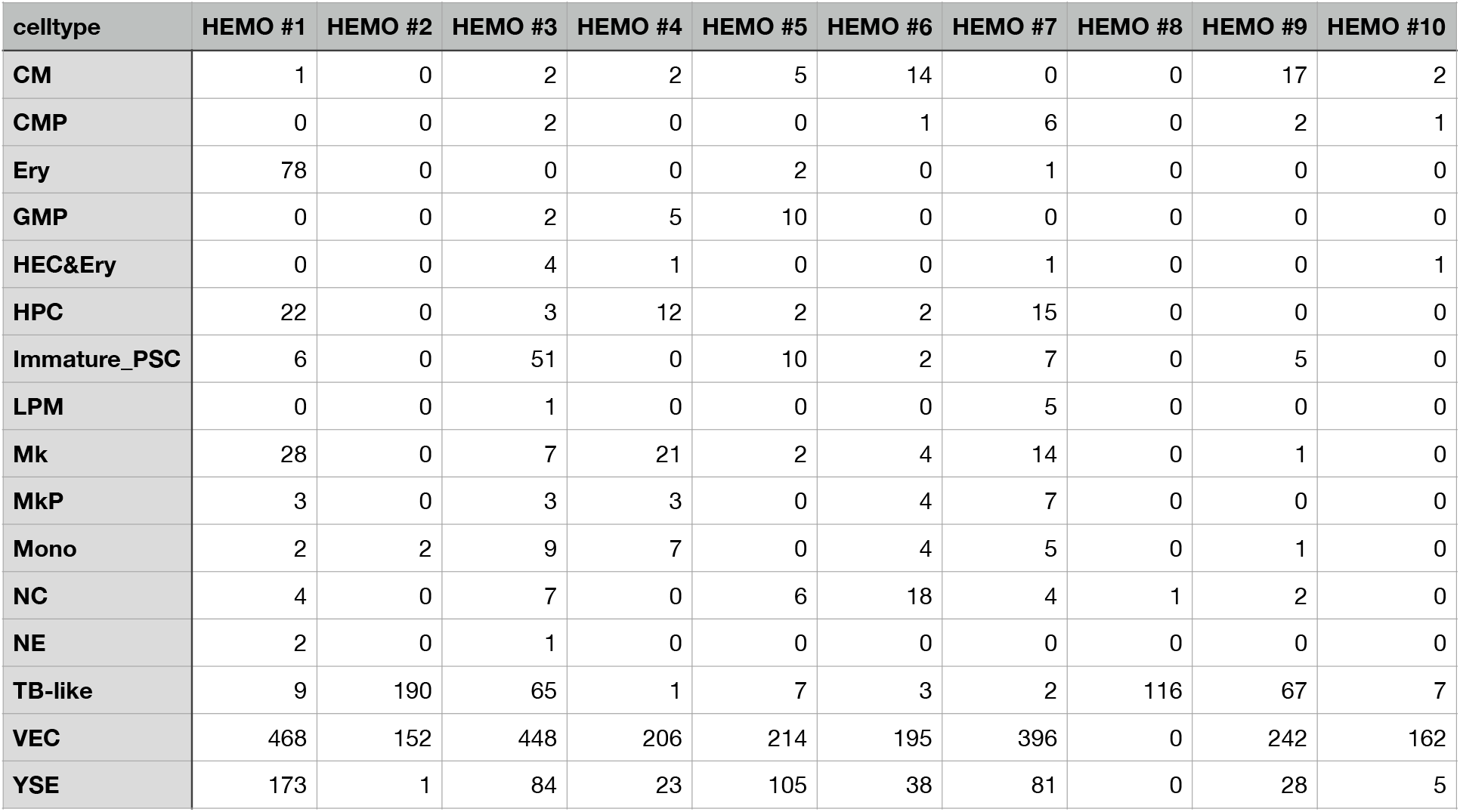
Cell proportion of HEMOs. Cell number of each HEMO imputed by SpatialScope based on scRNA-seq reference. Labels are correlated to Fig. 3C.

| celltype | HEMO #1 | HEMO #2 | HEMO #3 | HEMO #4 | HEMO #5 | HEMO #6 | HEMO #7 | HEMO #8 | HEMO #9 | HEMO #10 |
| --- | --- | --- | --- | --- | --- | --- | --- | --- | --- | --- |
| CM | 1 | 0 | 2 | 2 | 5 | 14 | 0 | 0 | 17 | 2 |
| CMP | 0 | 0 | 2 | 0 | 0 | 1 | 6 | 0 | 2 | 1 |
| Ery | 78 | 0 | 0 | 0 | 2 | 0 | 1 | 0 | 0 | 0 |
| GMP | 0 | 0 | 2 | 5 | 10 | 0 | 0 | 0 | 0 | 0 |
| HEC&Ery | 0 | 0 | 4 | 1 | 0 | 0 | 1 | 0 | 0 | 1 |
| HPC | 22 | 0 | 3 | 12 | 2 | 2 | 15 | 0 | 0 | 0 |
| Immature_PSC | 6 | 0 | 51 | 0 | 10 | 2 | 7 | 0 | 5 | 0 |
| LPM | 0 | 0 | 1 | 0 | 0 | 0 | 5 | 0 | 0 | 0 |
| Mk | 28 | 0 | 7 | 21 | 2 | 4 | 14 | 0 | 1 | 0 |
| MkP | 3 | 0 | 3 | 3 | 0 | 4 | 7 | 0 | 0 | 0 |
| Mono | 2 | 2 | 9 | 7 | 0 | 4 | 5 | 0 | 1 | 0 |
| NC | 4 | 0 | 7 | 0 | 6 | 18 | 4 | 1 | 2 | 0 |
| NE | 2 | 0 | 1 | 0 | 0 | 0 | 0 | 0 | 0 | 0 |
| TB-like | 9 | 190 | 65 | 1 | 7 | 3 | 2 | 116 | 67 | 7 |
| VEC | 468 | 152 | 448 | 206 | 214 | 195 | 396 | 0 | 242 | 162 |
| YSE | 173 | 1 | 84 | 23 | 105 | 38 | 81 | 0 | 28 | 5 |

